# Development and application of nbLIBRA-seq for high-throughput discovery of antigen-specific nanobodies

**DOI:** 10.64898/2025.12.30.697047

**Authors:** Sabina E. W. Leonard, Perry T. Wasdin, Katherine Webb, Parastoo Amlashi, Jeffrey C. Rathmell, Benjamin W. Spiller, Brian E. Wadzinski, Ivelin S. Georgiev, Kelsey Voss

## Abstract

Nanobodies are of high interest in many fields of medicine and biotechnology due to their high stability, tissue penetration, and engineering adaptability compared to monoclonal antibodies. However, nanobody discovery has been limited by technologies that rely on laborious library generation, panning, and clone screening techniques. Here, we demonstrate the successful adaptation of Linking B-Cell Receptor to Antigen Specificity through Sequencing (LIBRA-seq) to immunized alpacas for the rapid identification of antigen-specific nanobodies, derived from heavy-chain antibodies. We validated for nanobody discovery (nbLIBRA-seq) in two different disease settings. First, we identified over 300 antigen-specific heavy chain antibodies against human Transferrin Receptor (TfR1), also known as CD71, from a single alpaca blood sample. Experimental validation showed nbLIBRA-seq was able to identify nanobodies that exhibit specific binding to CD71, with two nanobodies also showing receptor internalization on human T cells. In a separate experiment, we tested the ability of nbLIBRA-seq to perform nanobody discovery with multiple antigens in the antigen screening library. Using fusion glycoproteins from the related respiratory syncytial virus (RSV) and human metapneumovirus (hMPV), 1,125 antigen-specific heavy-chain expressing B cells were recovered via nbLIBRA-seq. A subset of these nanobodies was validated experimentally to possess the target antigen specificity. Together, our results illustrate the potential of nbLIBRA-seq to rapidly identify antigen-specific heavy chain antibodies for a range of diverse targets, a capability that will be of critical significance for the effective and efficient development of novel nanobody-based therapeutics against targets of biomedical significance.

## INTRODUCTION

Nanobodies are single-domain antibodies derived from heavy-chain only antibodies, produced naturally in camelids and sharks^1,2^. Resembling the V_H_ domains of human immunoglobulins, nanobodies are the smallest known naturally occurring antigen binding fragment and have demonstrated impressive potential for a variety of indications in pre-clinical and clinical studies^3–7^. Compared to canonical monoclonal antibodies, nanobodies demonstrate higher stability, increased tissue penetration and improved hydrophilicity, while retaining high affinity to antigen—making them ideal biomolecules for therapeutic development^3,4,8–13^. Additionally, nanobodies have >80% sequence conservation with human immunoglobulin heavy chain sequences, conferring low immunogenicity profiles and supporting the feasibility of human administration^3,4,13,14^. A major obstacle in the development of nanobody biologics has been the limited capabilities of nanobody discovery technologies. In particular, nanobody discovery has relied largely on the development of nanobody display libraries, which traditionally screen against a single antigen target at a time and are slow to produce^9,15–17^.

Here, we present nbLIBRA-seq, an adaptation to Linking B-Cell Receptor to Antigen Specificity through Sequencing (LIBRA-seq) for the rapid identification of antigen-specific heavy chain antibodies from immunized alpacas. LIBRA-seq is a rapid, high-throughput multi-omic approach that enables the discovery of antigen-specific B cells for large libraries of protein antigens within a single sample, through biotinylation and DNA-barcoding of antigens of interest^18–21^. Adaptation of this technology for the discovery of heavy chain antibodies enables rapid identification of B cells with specificity for one or several antigens of interest simultaneously, improving upon current methods for nanobody discovery.

As an initial proof-of-concept, we applied nbLIBRA-seq to nanobody discovery for the human Transferrin Receptor (TfR1), also known as CD71. We previously demonstrated CD71 as a desirable target in the treatment of autoimmune diseases including systemic lupus erythematosus (SLE) due to its role in T cell iron metabolism^22^. Although expressed on all cell types, targeting CD71 with a monoclonal antibody selectively depleted pro-inflammatory T cells and spared regulatory T cells. Antibody binding resulted in receptor internalization without the uptake of iron, normalizing iron levels in SLE T cells. Therefore, a nanobody that binds and internalizes CD71 to lower iron levels would be ideal for therapeutic development^22^. CD71 nanobodies that cause receptor internalization are also a priority in development as drug delivery agents, especially across the blood brain barrier^23,24^.

A major advantage of LIBRA-seq over alternative antibody discovery technologies is the ability to simultaneously screen against multiple antigen targets. Such a capability can be critical in a variety of settings, including the targeted discovery of antibodies that can cross-react between multiple variants of the target antigen, assuring a lack of polyreactivity against unrelated antigens, cross-reactivity against orthologs from other species and efficient screening for multiple different targets from a single physical sample. To validate the ability of nbLIBRA-seq to simultaneously screen for nanobodies against multiple different targets, we applied this technology targeting respiratory syncytial virus (RSV) and human metapneumovirus (hMPV) fusion (F) glycoproteins as our antigens of interest. RSV and hMPV are related viral pathogens and two of the leading causes of respiratory tract infection globally^25–27^. Their F glycoproteins mediate viral fusion and are the primary target of neutralizing antibodies. Nanobodies are an especially desirable drug class for respiratory infections like RSV and hMPV, as they can be effectively administered as inhaled biologics directly to the viral niche^28,29^. RSV and hMPV nanobody discovery using nbLIBRA-seq showcases both the ability of this technology to simultaneously screen against multiple antigen targets and the potential for the discovery of new leads for further therapeutic development against these pathogens of high biomedical significance.

## RESULTS

### Adaptation of LIBRA-seq for Identification of Alpaca Heavy Chain Antibodies

For efficient antigen-specific nanobody discovery, we adapted LIBRA-seq for use with alpaca peripheral blood mononuclear cells (PBMCs) from immunized animals. LIBRA-seq is a high-throughput single-cell sequencing technology that can rapidly identify antigen-specific B cells through conjugation of antigens to unique oligonucleotide barcodes. We first designed a flow cytometry panel that enables the identification of antigen-bound IgG2 and IgG3 expressing B-cells (which produce heavy chain only antibodies). The final antibody panel for sorting of antigen-specific B cells was designed to isolate IgG+ (Subclasses 2+3 specific), VHH+ double-positive cells that were bound to streptavidin-BV421 and thus bound to our biotin-labeled antigen (**Figure 1A**). Prior to sorting, PBMCs from the CD71-immunized alpaca were incubated with labeled antigens to allow for B cell binding. Each antigen was labeled with a unique DNA barcode and biotinylated to allow for identification with streptavidin-BV421 staining. Briefly, DNA barcoding occurs through a stable, covalent reaction between S-Hynic labeled antigen and S-4FB labeled oligonucleotide. Our pilot panel included CD71 and two negative control antigens, BG505 T332N (HIV env) and CA/09 H1 NC99 (Influenza HA) (**Supplemental Figure 1A**). To obtain heavy chain B cell receptor sequences, target enrichment was performed following cDNA generation. The inner and outer alpaca-specific primers were generated based on previous work (Alpaca Outer Primer: 5’- GGTACGTGCTGTTGAACTGTTCC-3’, Alpaca Inner Primer #1: 5’- GATCACTAGTGGGGTCTTCGCTGTGGTGCG-3’, Alpaca Inner Primer #2: 5’- GATCACTAGTTTGTGGTTTTGGTGTCTTGGG-3’)^30,31^. The forward primers are not species-specific and were identical to the 10X-supplied primers used for BCR amplification (5’-GATCTACACTCTTTCCCTACACGACGC-3’) (**Figure 1B**). Following sequencing, the variable heavy and hinge regions of heavy-chain expressing antibodies are recovered, enabling us to determine isotype information in addition to antigen specificity. We refer to this technology for nanobody discovery as nbLIBRA-seq.

**Figure 1.**
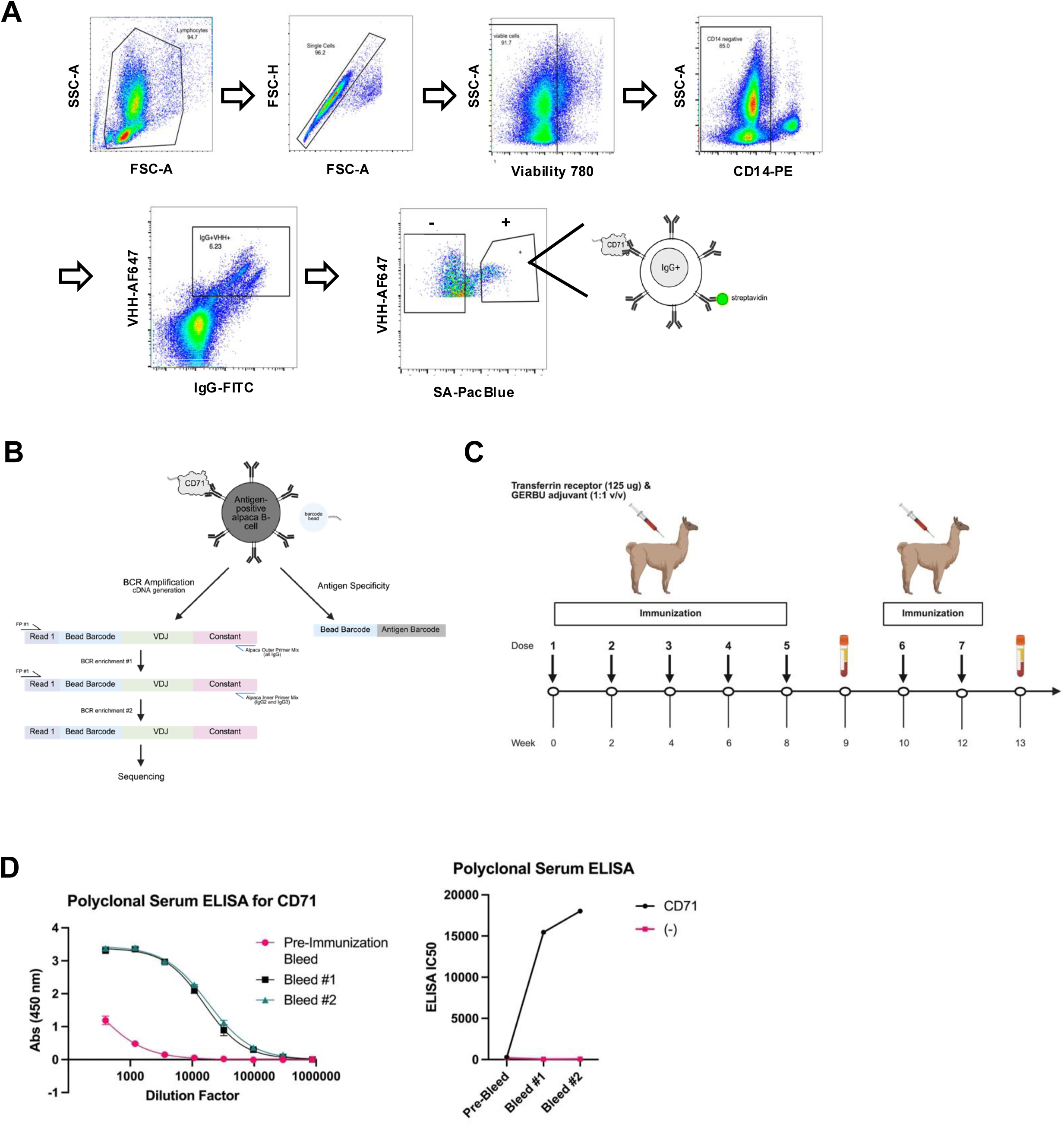
Alpaca immunization and development of LIBRA-seq with alpaca PBMCs. **(A)** LIBRA-seq sorting strategy to identify CD71-specific B cells. Alpaca PBMCs were incubated with biotinylated and oligo-barcoded human antigens, CD71 and negative controls, prior to sorting. CD14+ cells were excluded to remove monocytes. Cells positive for both IgG and VHH were gated for followed by antigen-bound B cells which were identified with fluorescently labeled streptavidin (SA). **(B)** LIBRA-seq workflow was adapted for use in alpacas by modifying BCR sequencing steps. Primers shown in blue indicated alpaca specific primers that anneal to the constant region of alpaca IgG genes. Antigen specificity was determined through computational deconvolution of barcodes. **(C)** Alpaca immunization schedule. One alpaca was immunized seven times at weeks 0, 2, 4, 6, 8,10 and 12. Blood was drawn at weeks 9 (Bleed #1) and 13 (Bleed #2). **(D)** Polyclonal serum ELISA from alpaca immunized with CD71. Serum pre-immunization, Bleed #1 (week 9) and Bleed #2 (week 13) were tested against CD71 and an unrelated antigen for binding. High binding was observed for CD71 following immunization while low binding was observed to CD71 prior to immunization and to an unrelated negative control antigen for all time points tested. Antibody binding ELISA titer of polyclonal serum from alpaca immunized with CD71 was calculated using a 4-parameter logistic curve fit to calculate the reciprocal serum dilution. ELISA was performed in technical duplicate and biological duplicate. Graph shown is a representative run.

### Elicitation of CD71-Specific Antibodies through Alpaca Immunization

To identify CD71-specific nanobodies, an alpaca was immunized seven times with the extracellular domain of recombinant human CD71 mixed 1:1 v/v with GERBU adjuvant. Blood was collected twice, after the fifth and seventh immunization, and used for isolation of peripheral blood mononuclear cells (PBMCs) and plasma (**Figure 1C**). Prior to nbLIBRA-seq, the plasma was tested for antibody binding and specificity to CD71. To do this, we conducted plasma ELISAs against CD71 and unrelated antigens (to which the alpaca is naïve) using pre-immunization blood, bleed #1, and bleed #2. Following immunization, there was greater antibody binding to CD71 while binding to an unrelated antigen was negligible at each time point (**Figure 1D**). Additionally, we performed dot blot analysis with plasma from pre-immunization and bleed #1 timepoints in the presence or absence of SDS (**Supplemental Figure 1B**). A reduction in antibody binding when treated with SDS indicated that some fraction of the B-cells are binding conformational epitopes of CD71. Taken together, these data illustrate that following immunization, a specific and robust immune response against CD71 was mounted.

### Discovery of CD71 Nanobodies Using nbLIBRA-seq

We obtained 362 antigen-specific B cells from our nbLIBRA-seq run after strict filtering criteria including removal of B cells with no antigen unique molecular identifier (UMI) counts, unproductive VDJ regions, and B cells with N>1 (which indicates more than 1 heavy chain). The B cells identified through nbLIBRA-seq used one of five distinct inferred germline genes and had CDR3 lengths between 5-23 amino acids (**Figures 2A+B**). Using calculated LIBRA-seq scores (LSS), we selected four lead nanobody candidates to express (**Figure 2C and Table S1**). The size and homogeneity of the nanobodies were evaluated by size exclusion chromatography and SDS-PAGE (**Supplemental Figure 2**). For all subsequent characterization, lead candidates were expressed as heavy chain antibodies (nanobody conjugated to human IgG1 Fc). To determine specific binding for CD71, ELISA was conducted with lead nanobody candidates and negative controls (**Figure 2D**). Lead candidates showed variable binding to CD71, with TFR1NBLS-01 and TFR1NbLS-02 showing the highest binding.

**Figure 2.**
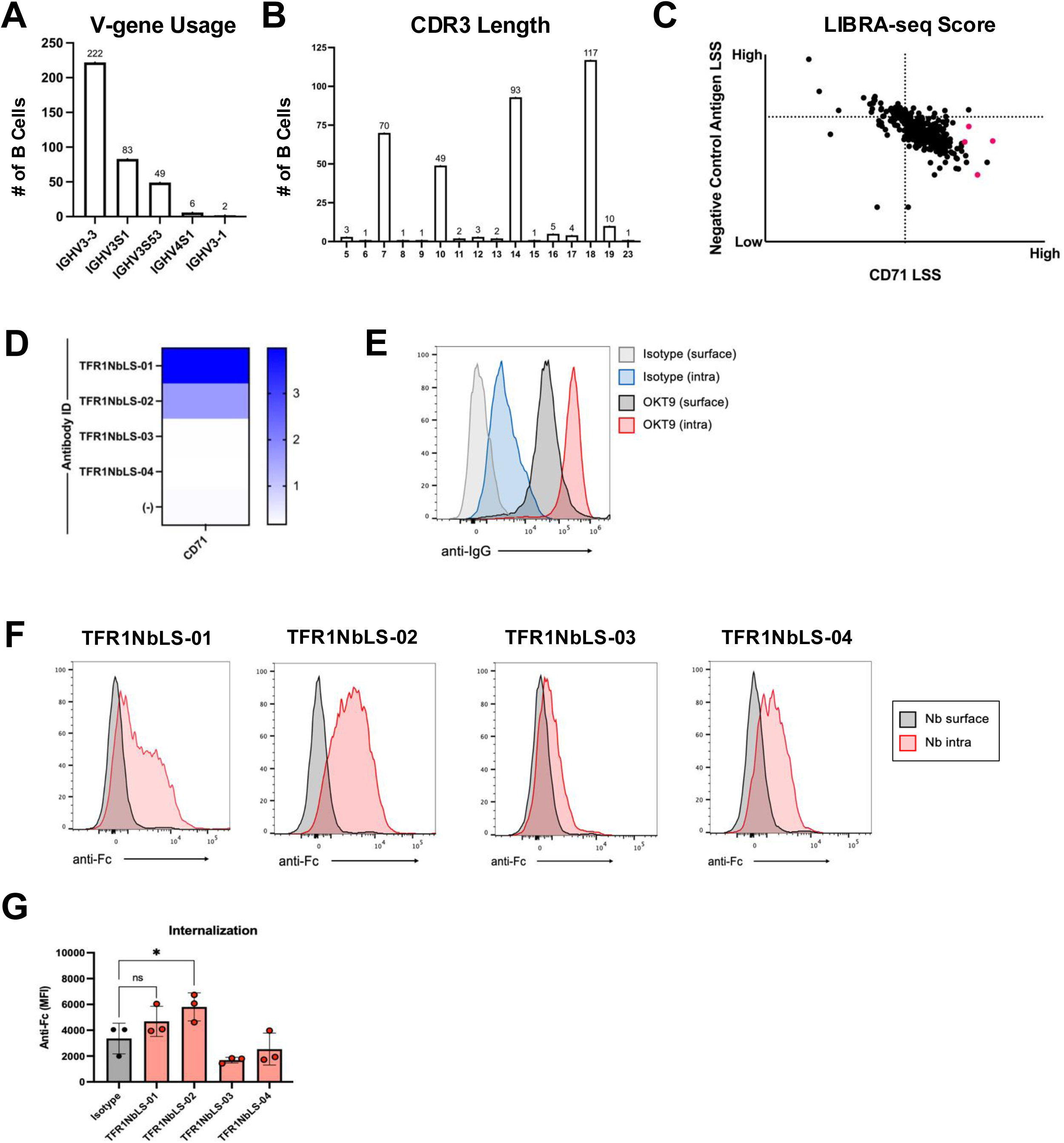
LIBRA-seq data following computational filtering. **(A)** V gene usage of antigen-specific heavy-chain antibodies. **(B)** CDR3 length of antigen-specific heavy-chain antibodies. **(C**) LIBRA-seq scores (LSS) of heavy-chain antibodies obtained from nbLIBRA-seq. Lead candidates, highlighted in pink and described in Table 1, were selected based off LIBRA-seq score (high LSS for CD71 and low LSS for negative control antigens). **(D)** ELISA area under the curve (AUC) values for lead nanobodies. Higher values indicate greater binding to respective antigen. **(E)** Assay for internalization of anti-CD71/TfR1 antibodies in Jurkat T cells. Surface-associated antibody versus intracellular (intra) was determined by differential flow cytometry staining permeabilizations with secondary antibody. **(F)** The assay in E was performed with nanobody candidates fused with an Fc domain to determine whether nanobodies were internalized or stay bound to the cell surface. **(G)** Quantification of receptor internalization from (F). One-way ANOVA.

To measure functional activity, candidate nanobodies were used in an TfR1 internalization assay adapted from our previous work, which showed the internalization of the anti-CD71 IgG antibody clone OKT9 after 30 minutes (**Figure 2E**). Using Jurkat human T cells, the nanobodies showed variable levels of TfR1 internalization, with TFR1NBLS-01 and TFR1NbLS-02 demonstrating the greatest amount of receptor internalization, with 1.39 and 1.73-fold increases in mean fluorescence intensity (MFI), respectively. (**Figure 2F+G**). Together, this work validates that LIBRA-seq identified nanobodies bind to their target antigen, CD71, and exhibit functionally relevant characteristics including TfR1 internalization.

### Application of nbLIBRA-seq for Simultaneous Discovery of Antigen-Specific Nanobodies against Multiple Different Targets

To further validate nbLIBRA-seq, we sought to screen for antigen-specific nanobodies in another alpaca immunized with different antigens – RSV F protein and hMPV F protein variants. Similar to the pilot run, plasma ELISA’s were conducted pre- and post-immunization against immunogens and an unrelated negative control antigen to ensure a robust and specific immune response was mounted (**Figure 3A**). To improve upon our true positivity rate, each antigen was labeled with two unique barcodes and only antigens positive for both barcodes were considered to have positive signal for that antigen (**Supplemental Figure 3**, **Supplemental Figure 4B**). Using the same cell-sorting and sequencing strategy as the CD71 nbLIBRA-seq run, this experiment recovered 3296 B cells. We then applied filtering criteria to remove B cells with no antigen UMI counts, unproductive/incomplete VDJ Regions and B cells with N>1 (indicating more than 1 HC per barcode). Following filtering, 716 B cells had positive signal for RSV F and 460 had positive signal for hMPV F (**Figure 3B**). The B cells identified through nbLIBRA-seq used one of 31 inferred germline V genes and had CDR3 lengths between 3-29 amino acids (**Figures 3C+D**). Of these, we selected 19 for experimental validation (**Figure 3B, Table S2**). In addition to selecting nanobodies with specificity for either RSV or hMPV, we also sought to determine if there are B cells that exhibited cross-reactivity between these two antigens. However, since there were no B cells with high-confidence signals for such cross reactivity, we elected to select B-cells with lower confidence signals (**Supplemental Figure 4A**). Leads were expressed as recombinant nanobody-Fc fusion (human IgG1) and assessed for binding via ELISA. Nanobodies screened bound their predicted antigen via ELISA (**Figure 3E**). Importantly, hMPV-reactive nanobodies bound wild-type hMPV F and hMPV F (D280N), a clinically relevant mutation that renders resistance to many published hMPV monoclonals. Successful identification and validation of heavy-chain only antibodies to viral glycoproteins in addition to previously studied autoantigens provides further validation that nbLIBRA-seq is a useful strategy to identify antigen-specific heavy chain antibodies and nanobodies for diverse antigens of interest.

**Figure 3.**
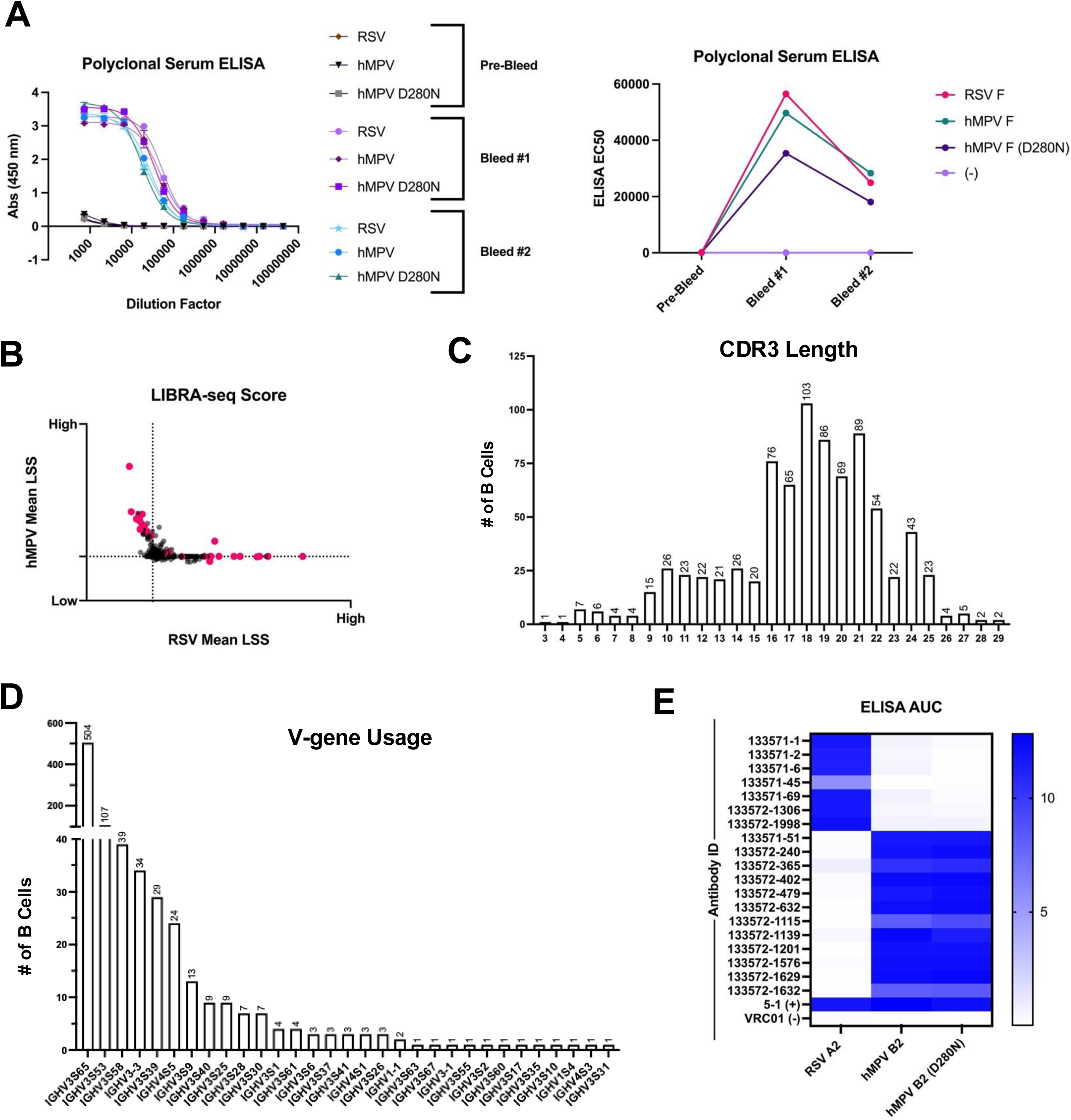
RSV and hMPV nanobodies. **(A)** Polyclonal serum ELISA from alpaca immunized with RSV and hMPV variants. Serum from pre-immunization, Bleed #1 (week 9) and Bleed #2 (week 13) were tested against immunogens and an unrelated antigen for binding. High binding was observed for immunogens following immunization while low binding was observed to immunogens prior to immunization and to an unrelated negative control antigen for all time points tested. Antibody binding ELISA titer of polyclonal serum was calculated using a 4-parameter logistic curve fit to calculate the reciprocal serum dilution. ELISA was performed in technical and biological duplicate. **(B)** LIBRA-seq scores (LSS) of heavy-chain antibodies obtained from nbLIBRA-seq. Lead candidates, highlighted in pink and described in Table 2, were selected based off LIBRA-seq score**. (C)** CDR3 length of antigen-specific heavy-chain antibodies. **(D)** V gene usage of antigen-specific heavy-chain antibodies. **(E)** ELISA area under the curve (AUC) values for lead nanobodies. Higher values indicate greater binding to respective antigen.

## DISCUSSION

Here we developed a novel method for the rapid identification of antigen-specific heavy chain antibodies from immunized alpacas. Using alpaca PBMCs, we identified CD71, RSV and hMPV specific heavy-chain antibodies and validated their characteristics through a variety of binding and functional assays. The nanobodies discovered reflected signatures of the camelid immunoglobulin repertoire including reduced V gene diversity and long CDR3 lengths^10^. Specifically, the majority of nanobodies we identified utilized the most reported IGHV germline genes – (IGHV3-3, IGHV3S53, IGHV3S61, IGHV3S65, or IGHV3S66) which suggests that the nanobodies discovered may be representative of the typical immunoglobulin repertoire^30,32^. Additionally, the median CDR3 length of the nanobodies discovered in the pilot CD71 nbLIBRA-seq run was 14 amino acids (mean 13.49 with standard deviation 4.31) while the median CDR3 length of nanobodies discovered in the RSV/hMPV nbLIBRA-seq run was 18 (mean 17.95 with standard deviation 4.42)^33^. The longer average CDR3 length of nanobodies compared to conventional antibodies may enable access to discrete epitopes inaccessible to larger antibodies, providing novel mechanisms of binding^34–36^.

Compared to previously published anti-CD71 nanobodies, ours are the first discovered without using phage-display technology and exhibit similar affinity to antigen^24,37,38^. Further, the use of nbLIBRA-seq enables the identification of hundreds of B cells simultaneously with antigen-binding predictions to both desired and negative-control antigens, which is unique to nbLIBRA-seq. Additionally, the hMPV and RSV nanobodies discovered in our nbLIBRA-seq run demonstrate significant genetic diversity. The ability to screen against multiple antigens simultaneously (e.g., hMPV +/− D280N mutation) enables the selection of nanobodies that demonstrate breadth. This is of key importance for viral targeting nanobodies as viruses often acquire mutations (like D280N) that render current therapeutics ineffective.

While the nanobodies we described may also be of interest for further preclinical assessment and development, the primary focus of the efforts described here is to perform proof-of-concept studies for the validation of nbLIBRA-seq as a novel, high-throughput tool that can be applied to discover nanobodies for any antigen of interest. This represents a large technical advance as it enables screening of several antigens of interest simultaneously which is important for several reasons, including the identification of cross-reactive nanobodies. Further, the implementation of nbLIBRA-seq enables various engineering efforts, including the development of antibody-drug conjugates and creation of multi-specific antibody constructs. nbLIBRA-seq can expedite these efforts as a single run can yield hundreds of antigen-specific nanobodies with variable functional characteristics. Together, these results establish nbLIBRA-seq as a versatile strategy for quickly discovering antigen-specific heavy chain antibodies, an advance that holds broad significance for nanobody engineering and therapeutic design.

## RESOURCE AVAILABILITY

### Lead Contact

Further information and requests for resources and reagents should be directed to and will be fulfilled by the Lead Contact, Dr. Kelsey Voss.

## ACKNOWLEDGEMENTS

We thank all members of the Georgiev laboratory for their support and feedback. We thank David Flaherty, Olivia X. Murfield, Emma McLaughlin, and Brittany Matlock from the VUMC Flow Cytometry Shared Resource for their help with cell sorting. Flow cytometry experiments were performed in the VUMC Flow Cytometry Shared Resource which is supported by the Vanderbilt Ingram Cancer Center (P30 CA68485) and the Vanderbilt Digestive Disease Research Center (DK058404). We thank Angela Jones, Jamie Roberson, and Latha Raju with the Vanderbilt Technologies for Advanced Genomics Core (VANTAGE) for providing technical assistance with library production and sequencing. VANTAGE is supported in part by CTSA (5UL1 RR024975-03), the Vanderbilt-Ingram Cancer Center (P30 CA068485), the Vanderbilt Vision Center (P30 EY08126), and NIH/NCRR (G20 RR030956). This research was funded, in part, by the Vanderbilt Center for Immunobiology (VCI), the Advanced Research Projects Agency for Health (ARPA-H 1AY2AX000077) and NIH R01 AI175245. The funders had no role in the conceptualization or execution of any studies or drafting of the manuscript. The views and conclusions contained in this document are those of the authors and should not be interpreted as representing the official policies, either expressed or implied, of the US government.

## AUTHOR CONTRIBUTIONS

Conceptualization and Methodology: S.E.W.L., I.S.G., K.V.; Investigation: S.E.W.L., P.W., K.W., P.A., B.S., B.W., K.V.; Writing – Original Draft: S.E.W.L., I.S.G., K.V.; Writing – Reviewing & Editing: All authors; Funding Acquisition: J.C.R., I.S.G., K.V.; Resources: B.S., B.W., J.C.R., K.V., I.S.G., K.V.; Supervision: B.S., B.W., J.C.R., I.S.G., K.V.

## METHODS

### Alpaca Immunization

Two alpacas (females) were used for CD71 and RSV/hMPV immunization. All immunizations were conducted by Turkey Creek Biotechnology (Waverly, TN) in accordance with IACUC protocol 18-02. Immunizations were conducted by Turkey Creek Biotechnology (Waverly, TN) in accordance with IACUC protocol 18-02. Briefly, an alpaca (female) was immunized 7 times subcutaneously approximately every 2 weeks with 150-300 ug of purified transferrin receptor in Gerbu adjuvant (1:1 v/v). Blood was drawn one-week after the 5^th^ and 7^th^ injections, and peripheral blood mononuclear cells (PBMCs) were isolated from blood by Ficoll-paque density gradient centrifugation using SepMate centrifugal devices according to the manufacturer’s protocol (StemCell Technologies). PBMCs were slowly frozen to −80°C and transferred to liquid nitrogen for long-term storage until use. A second alpaca (female) was immunized 6 times with 150 ug recombinant RSV F protein and 150 ug hMPV F protein (immunization 1-3: D280 variant, immunization 4-6: N280 variant) every two weeks. Blood was drawn one week after the 4^th^ and 6^th^ immunization. Peripheral blood mononuclear cells (PBMCs) were isolated from blood by Ficoll-paque density gradient centrifugation using SepMate centrifugal devices according to the manufacturer’s protocol (StemCell Technologies). PBMCs were slowly frozen to −80°C and transferred to liquid nitrogen for long-term storage until use.

### Immunogen expression and purification

The immunogens used for immunization, LIBRA-seq and subsequent nanobody analysis were produced in Expi293F cells by transient transfection in FreeStyle F17 expression media (ThermoFisher) supplemented to a final concentration of 0.1% Pluronic Acid F-68 and 20% 4 mM L-glutamine using Expifectamine transfection reagent (ThermoFisher). Cells were cultured for 5-6 days at 8% CO_2_ saturation and 37°C with shaking. Following transfection, cultures were centrifuged at 4000 rcf for 20 minutes and supernatant was filtered with Nalgene Rapid Flow Disposable Filter Units with PES membrane (0.45 mm) then run slowly over an affinity column (HisTrap HP followed by StrepTrap XT) at 4°C. The protein elution was buffer-exchanged 3 times into PBS and concentrated using a 30 kDa Amicon Ultra centrifugal filter unit. Concentrated protein was run on a Superose 6 Increase 10/300 GL or Superdex 200 Increase 10/200 GL on the AKTA FPLC system. Peaks corresponding to properly formed species were identified based on elution volume and visualized with SDS-PAGE. All proteins were quantified using UV/vis spectroscopy. Antigenicity of proteins was characterized by ELISA with known monoclonal antibodies specific for that antigen. Proteins were flash frozen and stored at −80°C until use.

### Biotinylation of antigens

Protein antigens were biotinylated using EZ-link Sulfo-NHS-Biotin No-Weigh kit (ThermoFisher) according to manufacturer’s instructions. A 50:1 biotin-to-protein molar ratio was used for all reactions. Transferrin receptor was biotinylated using BirA enzyme which enables site specific biotinylation at the AviTag sequence.

### Oligonucleotide barcodes

We used oligos that possess a 15 bp antigen barcode, a sequence capable of annealing to the template switch oligo that is part of the 10X bead-delivered oligos and contain truncated TruSeq small RNA read 1 sequences in the following structure: 5’-CCTTGGCACCCGAGAATTCCANNNNNNNNNNNNNNNCCCATATAAGA*A*A-3’, where Ns represent the antigen barcode. Oligos were ordered from Sigma-Aldrich and IDT with a 5’ amino modification and HPLC purified. The following antigen barcodes were used: TCCTTTCCTGATAGG (BG505), TACGCCTATAACTTG (NC99), and CAGATGATCCACCAT (CD71). For the secondary validation of nanobody LIBRA-seq, each antigen was labeled with two unique barcodes. For each antigen, we used the following sequences: AATACCTCACAACGT (RSV), GGACGTGCACACTGA (RSV), CGCGTGTACCGGCAT (hMPV B2), TAGTGTCACTATGTT (hMPV B2), CCCGCTCCACAGCAA (hMPV B2 D280N), AAGTT TACCCTTCA (hMPV B2 D280N), GTTTCATAGACGCCC (BG505), and TGATAGTCGATAACC (BG505).

### Conjugation of oligonucleotide barcodes to antigens

For each antigen, a unique DNA barcode was directly conjugated to the antigen using a SoluLINK Protein-Oligonucleotide Conjugation kit (TriLink, S-9011) according to the manufacturer’s protocol. Briefly, the oligo and protein were desalted, and then the amino-oligo was modified with the 4FB crosslinker, and the biotinylated antigen protein was modified with S-HyNic. Next, the 4FB-oligo and the HyNic-antigen were mixed. This causes a covalent bond to form between the protein and the oligonucleotide. The concentration of the antigen-oligo conjugates was determined by a BCA assay, and the HyNic molar substitution ratio of the antigen-oligo conjugates was analyzed using the NanoDrop according to the Solulink protocol guidelines. Excess oligonucleotide was removed from the protein-oligonucleotide conjugates using Superdex 200 Increase 10/300 GL or Superose 6 Increase10/300 GL on AKTA FPLC. Protein-oligonucleotide conjugates were verified with SDS-PAGE and silver stain. The optimal amount of antigen-oligonucleotide conjugates to use in antigen-specific B cell sorting was determined through titrations on Ramos B Cells with known BCR specificity.

### Enrichment of antigen-specific alpaca B cells

Frozen alpaca PBMCs (isolated from bleed #1 post immunization) were thawed and resuspended in 10 ml of pre-warmed RPMI 1640 medium supplemented with 10% heat-inactivated Fetal bovine serum (FBS) (cRPMI). The cells were then washed with ten milliliters of cRPMI at 300 rcf for 5 minutes. Cells were counted and viability was determined using Trypan Blue. Cells were resuspended in 2 ml of phenol red-free RPMI supplemented with 0.5% rat serum (staining buffer) and stained with cell markers including viability dye (Ghost Red 780), PE-conjugated CD14 monoclonal antibody (TuK4), FITC-Conjugated AffiniPure Goat Anti-Alpaca IgG (Subclasses 2+3 specific), and AF647-Conjugated AffiniPure Goat Anti-Alpaca IgG (VHH domain) in the dark for 30 minutes at 4°C. Cells were then washed three times with staining buffer at 300 rcf for 5 minutes. Antigen-oligo conjugates were then added and incubated with the cells for 30 minutes in the dark at 4°C. Cells were then washed three times with staining buffer at 300 rcf for 5 minutes. Streptavidin-BV421 was added to label the cells with bound antigen and incubated for 15 minutes in the dark at 4°C. Cells were then washed three times with staining buffer, resuspended in staining buffer, and then sorted by FACS by the Vanderbilt Flow Cytometry Shared Resource core on a 5-laser FACS Aria III. Antigen-positive B cells were delivered to Vanderbilt Technologies for Advanced Genomics (VANTAGE) sequencing core at an appropriate target concentration for 10X Genomics library preparation and sequencing. FACS data was analyzed using FlowJo.

### 10X#Genomics single cell processing and next generation sequencing

Single-cell suspensions were loaded onto the Chromium Controller microfluidics device (10X Genomics) and processed using previously described methods for LIBRA-seq, with the exception that a custom primer set specific to alpaca was used for target enrichment steps^18,39^. Target enrichment step 1 consisted of a mix of Outer RP sequences 1-2 (final concentrations of 0.5mM each) and FP1 (final concentration 1mM). Target enrichment step 2 consisted of a mix of inner RP sequences 1-2 (final concentration of 0.5 mM each) and FP2 (final concentration of 1mM)^40^. A target capture of 10,000 B cells was used per 1/8 10X cassette for B cells. Slight modifications were made to intercept, amplify, and purify the antigen barcode libraries as previously described.

### Sequence processing and bioinformatics analysis

We followed our established pipeline, which takes paired-end FASTQ files of oligonucleotide libraires as input, to process and annotate reads for cell barcodes, unique molecular identifiers (UMIs) and antigen barcodes, resulting in a cell barcode-antigen barcode UMI count matrix^18^. B cell receptor contigs were processed Cell Ranger V(D)J v7.1.0 (10X Genomics) with a custom V(D)J reference. This reference was constructed by querying the IMGT/GENE-DB database for all *Vicugna pacos* IG gene sequences, which were then formatted using the Cell Ranger *mkref* pieline^41^. The antigen barcode libraries were processed using CellRanger count (10X Genomics). The cell barcodes that overlapped between the two libraries formed the basis of the subsequent analysis. Cell barcodes that only had non-functional heavy chain sequences as well as cells with multiple functional heavy chain sequences were eliminated, as these may represent multiplets. We also aligned the B cell receptor contigs (filtered_contigs.fasta file output by CellRanger, 10X Genomics) to IMGT references genes for sequence annotation using IMGT/HighV-Quest^42^. Finally, we determined the LIBRA-seq score for each antigen in the library for every cell as previously described. For alpaca gene assignments of the antibodies shown in figure, the immunoglobulin amino acid sequences for alpacas were obtained from and aligned to amino acid sequences of antibodies discovered in our work described here using Clustal Omega.

### Alpaca plasma ELISAs

Polyclonal plasma ELISA for specificity and magnitude of alpaca plasma response was conducted with plasma pre- and post-immunization. CD71 or an unrelated negative control immunogen (HIV Envelope) was plated at 2 ug/mL overnight at 4°C. The next day, plates were washed three times with PBS supplemented with 0.01% Tween20 (PBS-T) and blocked with 5% BSA in PBS-T. Plates were incubated at room temperature for two hours and then washed three times with PBS-T. Alpaca plasma was diluted starting at 1:400 (highest concentration) in 1% BSA in PBS-T, and then 3-fold down for a total of seven dilutions. Plates were incubated at room temperature for one hour and then washed three times with PBS-T. The secondary antibody, goat anti-alpaca conjugated to peroxidase, was added at 1:10,000 dilution in 1% BSA in PBS-T to the plates and incubated for one hour at room temperature. Plates were washed three times and then developed by adding TMB substrate to each well. The plates were incubated at room temperature for ten minutes, and then 1N sulfuric acid was added to stop the reaction. Plates were read at 450 nm. ELISAs were repeated in technical and biological duplicate. ELISA IC50 was calculated from interpolation of Sigmoidal, 4PL, X is concentration fit. Polyclonal plasma ELISAs were also conducted with plasma from the RSV/hMPV-immunized alpaca as described above.

### Dot blot assays

One microliter of 0.1, 0.01, 0.001, and 0.0001 mg/mL stock solutions of transferrin receptor in SDS sample buffer or PBS were spotted on Amersham Protran Supported 0.2 mm Nitrocellulose (GE Healthcare Life Sciences). The membranes were allowed to dry, wetted in Tris-buffered saline (TBS), blocked in Intercept Blocking Buffer (LI-COR), and then incubated overnight with alpaca plasma (from immunized animal and control animal) diluted 1:200 in Intercept Blocking Buffer. After washing with TBS/0.1% Tween, the membranes were incubated with an 800 fluorophore-labeled goat anti-alpaca secondary antibody, washed with TBS/0.1% Tween, and then imaged on the Odyssey Infrared Imaging system (LI-COR).

### Nanobody purifications

For each nanobody, the variable heavy gene was synthesized as cDNA and inserted into pcDNA3.1(+). Nanobodies were expressed with Expifectamine transfection reagent (Thermo Fisher Scientific) in Expi293F cells in FreeStyle F17 expression media (Thermo Fisher Scientific) supplemented with 0.1% Pluronic Acid F-68 and 20% 4 mM L-glutamine. Cells were cultured for 5 days at 8% CO_2_ saturation and 37°C with shaking. Five days post transfection, cells were collected and centrifuged at 4000 rpm for 20 minutes. Supernatant was filtered with Nalgene Rapid Flow Disposable Filter Units with PES membrane (0.22 μm) and purified with affinity (StrepTrap XT and HisTrap HP) chromatography at 4°C. The protein elution was buffer exchanged 3 times into PBS and concentrated using a 3 kDa Amicon Ultra centrifugal filter unit. Concentrated protein was run on a Superdex 75 Increase 10/300 GL on the AKTA FPLC system. Peaks corresponding to properly formed species were identified based on elution volume and visualized with SDS-PAGE. All proteins were quantified using UV/vis spectroscopy.

### Receptor internalization assay

Jurkat T cells were maintained in cRPMI (10% HI FBS + 1% anti-anti). Roughly 500,000 cells were transferred to a 5ml FACS tube and washed with 2 ml 1X PBS. Cells were starved for 1 hour in 1 ml of PBS + 2% BSA at 37°C and then incubated with 2 ug/ml of each nanobody or antibody. After 30 minutes, 2 ml of ice cold FACS buffer (PBS + 2% HI FBS + 2 mM EDTA) was added, and cells were washed with an additional 2 ml of FACS buffer. For surface staining, cells were then incubated with fixable viability dye and secondary antibody or isotype control. For intracellular staining of nanobodies, surface staining only consisted of the viability dye. After washing with 2 ml of FACS buffer, cells were then treated with BD Cytofix/Cytoperm buffer for 30 minutes on ice and washed again. Intracellular antibody staining was performed with intracellular staining perm wash buffer (Biolegend 421002). Antibodies for nanobody detection: BV421 anti-human IgG Fc (Biolegend 410703) or isotype control BV421 Rat IgG2a (Biolegend 400549) both used at 1:500. Antibody for OKT9 detection: PE rat anti-mouse IgG1 (BD 550083). Cells were resuspended in FACS buffer and analyzed using the Cytek Aurora flow cytometer.

### Nanobody-FC ELISA

ELISAs were conducted to evaluate the binding of LIBRA-seq identified nanobodies to their predicted antigen. Soluble, purified antigen was plated at a concentration of 2 ug/mL overnight at 4°C. The next day, plates were washed three times with PBS supplemented with 0.01% Tween20 (PBS-T) and blocked with 5% BSA in PBS-T. Plates were incubated at room temperature for two hours and then washed three times with PBS-T. Primary antibodies were diluted in 1% BSA in PBS-T, starting at a concentration of 10 μg/mL with a serial 1:5 dilution, then added to the plate. Plates were incubated for an hour at room temperature and then washed three times with PBS-T. The secondary antibody, goat anti-human IgG conjugated to peroxidase, was added at a dilution of 1:10,0000 in 1% BSA in PBS-T to the plates and then incubated for one hour at room temperature. Plates were washed three times with PBS-T and developed with the addition of TMB substrates. Plates were incubated at room-temperature in the dark for 10 minutes before the reaction was stopped with 1N sulfuric acid. Plates were read at 450 nm. Data shown is one biological replicate. Each ELISA was repeated in biological and technical duplicate. The area under the curve (AUC) was calculated using GraphPad Prism Version 10.5.0.

### Quantification and Statistical Analysis

All quantification and statistical analyses were performed using GraphPad Prism 10.5.0. software. Repeated measures one-way ANOVA was performed for comparisons. p < 0.05 was considered as statistically significant with mean ± SD. All statistical details of experiments can be found in the figure legends.

