## Supplemental Materials for "Development and application of nbLIBRA-seq for high-throughput discovery of antigen-specific nanobodies"

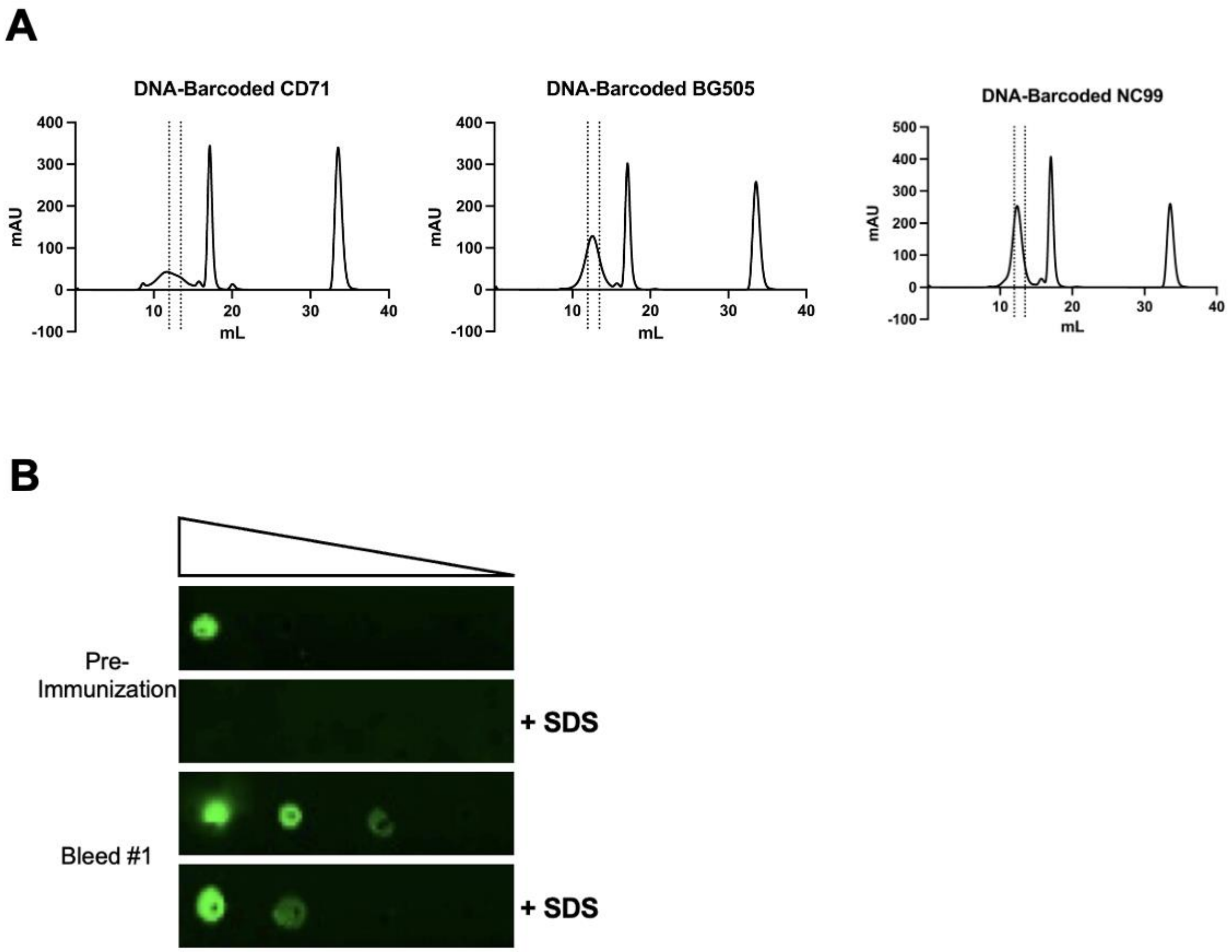


**Supplementary Figure 1.** Antigens used for LIBRA-seq analysis. **(A)** Size exclusion chromatograms of labeled antigens used in LIBRA-seq. Antigens were expressed recombinantly in mammalian cells and purified with affinity and size exclusion columns. Dashed lines indicate fractions collected indicative of DNA-barcoded and biotinylated antigens. **(B)** Dot blot depicting serum response to CD71 pre-immunization and Bleed #1. SDS disrupts disulfide bonds. Reduced serum response in presence of SDS indicates some amount of serum antibodies recognize conformational epitopes.

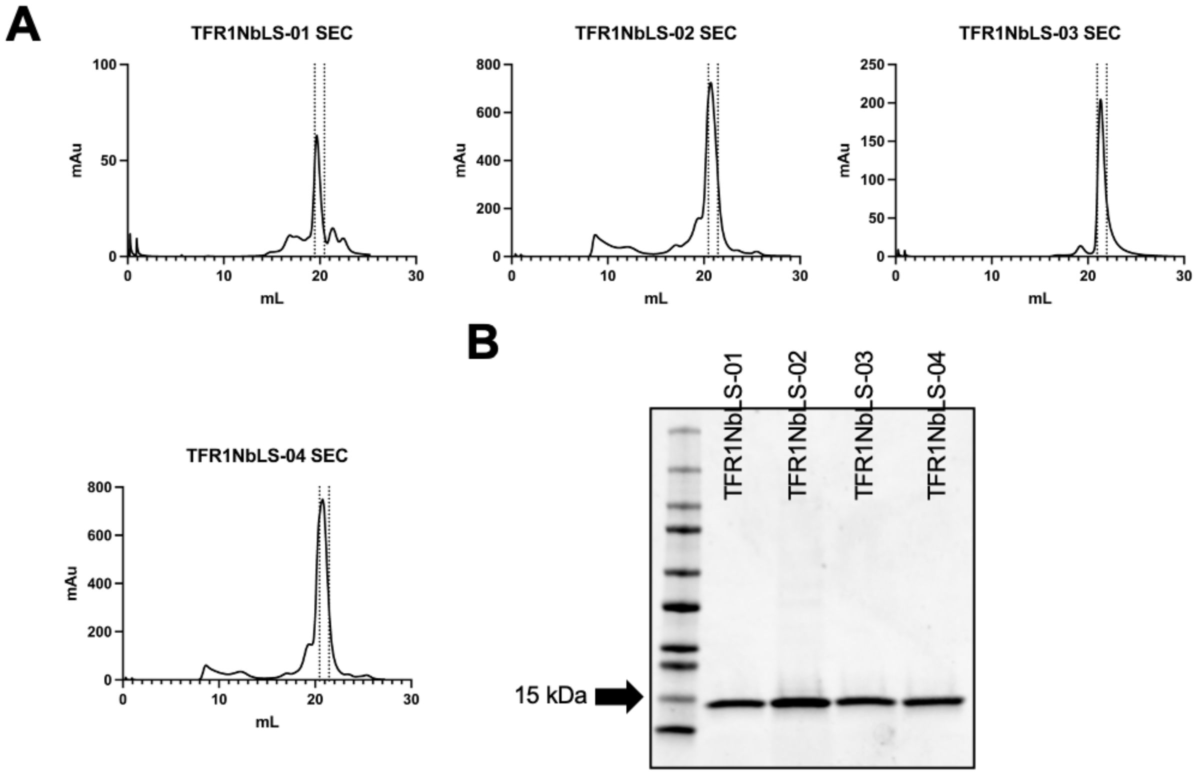


**Supplementary Figure 2. (A)** Size exclusion chromatograms of lead nanobodies identified from nbLIBRA-seq. Nanobodies were expressed recombinantly in mammalian cells and purified with affinity and size exclusion columns. Dashed lines indicate fractions collected indicative of nanobodies expected size. **(B**) SDS-PAGE gel of purified nanobodies from (A). Nanobodies have an expected size of 15 kDa.

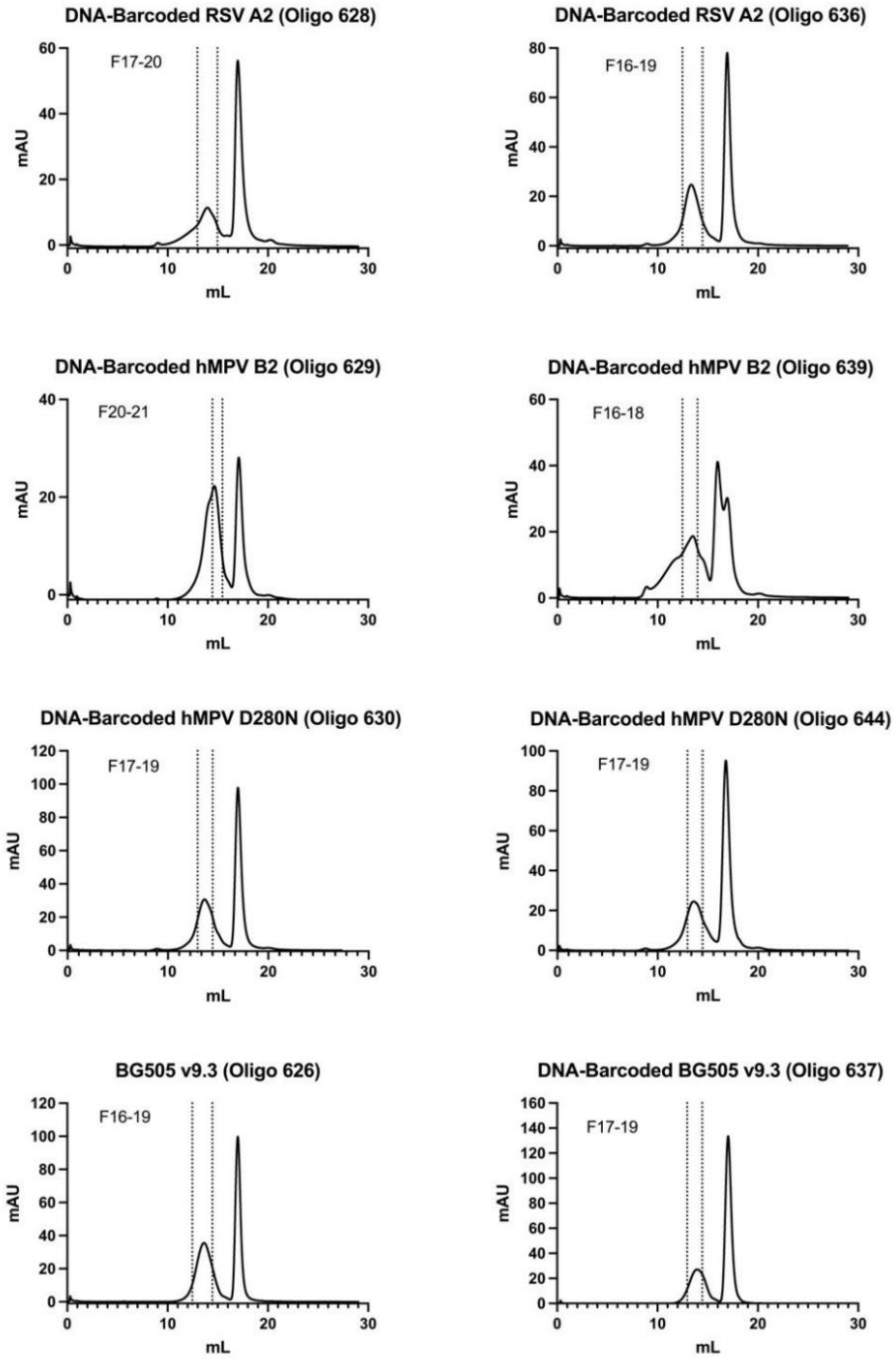


**Supplementary Figure 3.** Antigens used for nbLIBRA-seq analysis. Size exclusion chromatograms of labeled antigens used in nbLIBRA-seq. Antigens were expressed recombinantly in mammalian cells and purified with affinity and size exclusion columns. Antigens were labeled with two unique barcodes and purified separately. Dashed lines indicate fractions collected indicative of DNA-barcoded and biotinylated antigens.

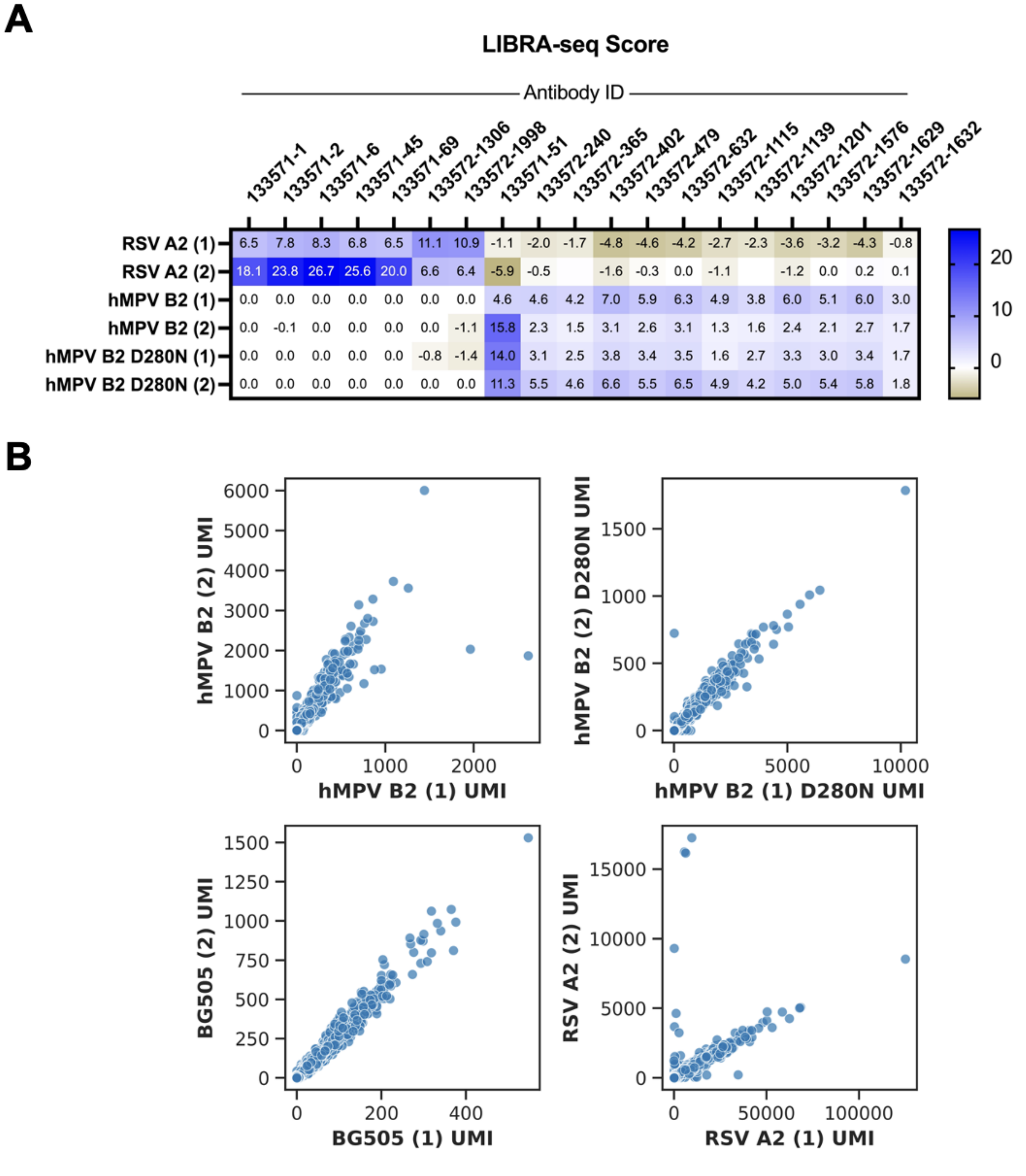


| | 133571-1 | 133571-2 | 133571-6 | 133571-45 | 133571-69 | 133572-1306 | 133572-1998 | 133571-51 | 133572-240 | 133572-365 | 133572-402 | 133572-479 | 133572-632 | 133572-1115 | 133572-1139 | 133572-1201 | 133572-1576 | 133572-1629 | 133572-1632 |
|---|---|---|---|---|---|---|---|---|---|---|---|---|---|---|---|---|---|---|---|
| RSV A2 (1) | 6.5 | 7.8 | 8.3 | 6.8 | 6.5 | 11.1 | 10.9 | -1.1 | -2.0 | -1.7 | -4.8 | -4.6 | -4.2 | -2.7 | -2.3 | -3.6 | -3.2 | -4.3 | -0.8 |
| RSV A2 (2) | 18.1 | 23.8 | 26.7 | 25.6 | 20.0 | 6.6 | 6.4 | -5.9 | -0.5 | | -1.6 | -0.3 | 0.0 | -1.1 | | -1.2 | 0.0 | 0.2 | 0.1 |
| hMPV B2 (1) | 0.0 | 0.0 | 0.0 | 0.0 | 0.0 | 0.0 | 0.0 | 4.6 | 4.6 | 4.2 | 7.0 | 5.9 | 6.3 | 4.9 | 3.8 | 6.0 | 5.1 | 6.0 | 3.0 |
| hMPV B2 (2) | 0.0 | -0.1 | 0.0 | 0.0 | 0.0 | 0.0 | -1.1 | 15.8 | 2.3 | 1.5 | 3.1 | 2.6 | 3.1 | 1.3 | 1.6 | 2.4 | 2.1 | 2.7 | 1.7 |
| hMPV B2 D280N (1) | 0.0 | 0.0 | 0.0 | 0.0 | 0.0 | -0.8 | -1.4 | 14.0 | 3.1 | 2.5 | 3.8 | 3.4 | 3.5 | 1.6 | 2.7 | 3.3 | 3.0 | 3.4 | 1.7 |
| hMPV B2 D280N (2) | 0.0 | 0.0 | 0.0 | 0.0 | 0.0 | 0.0 | 0.0 | 11.3 | 5.5 | 4.6 | 6.6 | 5.5 | 6.5 | 4.9 | 4.2 | 5.0 | 5.4 | 5.8 | 1.8 |

**Supplementary Figure 4.** **(A)** LIBRA-seq scores for lead nanobody candidates. **(B)** UMI counts for each antigen barcode. Each point represents a heavy chain-expressing B cell and the UMI for each distinct barcode within a given antigen. Higher UMI values indicate a greater amount of antigen bound to that B cell.

**Supplementary Table 1:**

| Name | V Gene | D Gene | J Gene | V Identity (%) | CDR3 Length (AA) |
|---|---|---|---|---|---|
| TFR1NbLS-01 | IGHV3-3 | IGHD1 | IGHJ4 | 86 | 17 |
| TFR1NbLS-02 | IGHV4S1 | IGHD2 | IGHJ4 | 86 | 19 |
| TFR1NbLS-03 | IGHV3S1 | IGHD5 | IGHJ4 | 80 | 7 |
| TFR1NbLS-04 | IGHV3-3 | IGHD3 | IGHJ4 | 68 | 18 |

**Supplementary Table 1.** Lead candidates chosen from CD71 LIBRA-seq analysis and V-, D-, J- inferred gene usage, percent V gene identity to germline and CDR3 length.

**Supplementary Table 2:**

| Name | V Gene | D Gene | J Gene | V Identity (%) | CDR3 Length (AA) |
|---|---|---|---|---|---|
| 133571-1 | IGHV3S65 | ~ | IGHJ7 | 98.6 | 18 |
| 133571-2 | IGHV3S65 | ~ | IGHJ7 | 98.61 | 18 |
| 133571-6 | IGHV3S65 | ~ | IGHJ4 | 94.44 | 24 |
| 133571-45 | IGHV3S39 | ~ | IGHJ4 | 80.9 | 12 |
| 133571-69 | IGHV3S65 | ~ | IGHJ7 | 97.22 | 18 |
| 133572-1306 | IGHV3S53 | IGHD5 | IGHJ4 or IGHJ6 | 89.12 | 12 |
| 133572-1998 | IGHV3S53 | IGHD5 | IGHJ6 | 91.58 | 11 |
| 133571-51 | IGHV3S65 | ~ | IGHJ7 | 98.26 | 24 |
| 133572-240 | IGHV3S65 | ~ | IGHJ4 | 88.19 | 21 |
| 133572-365 | IGHV3S65 | ~ | IGHJ4 | 88.19 | 25 |
| 133572-402 | IGHV3S53 | IGHD2 | IGHJ6 | 95.09 | 16 |
| 133572-479 | IGHV3S65 | ~ | IGHJ4 | 88.54 | 21 |
| 133572-632 | IGHV3S53 | IGHD2 | IGHJ6 | 96.84 | 16 |
| 133572-1115 | IGHV3S65 | ~ | IGHJ4 | 87.85 | 21 |
| 133572-1139 | IGHV3S65 | ~ | IGHJ4 | 97.22 | 20 |
| 133572-1201 | IGHV3S65 | ~ | IGHJ4 | 87.15 | 21 |
| 133572-1576 | IGHV3S65 | ~ | IGHJ4 | 86.46 | 21 |
| 133572-1629 | IGHV3S53 | IGHD2 | IGHJ6 | 94.39 | 16 |
| 133572-1632 | IGHV3S65 | ~ | IGHJ7 | 99.31 | 23 |

**Supplementary Table 2**. Lead candidates chosen from RSV and hMPV LIBRA-seq analysis and V-, D-, J- inferred gene usage, percent V gene identity to germline and CDR3 length. ~ indicates ambiguous germline D gene assignment.
